# Monitoring single-cell dynamics of entry into quiescence during an unperturbed lifecycle

**DOI:** 10.1101/2020.11.25.395608

**Authors:** Basile Jacquel, Théo Aspert, Damien Laporte, Isabelle Sagot, Gilles Charvin

## Abstract

The life cycle of microorganisms is associated with dynamic metabolic transitions and complex cellular responses. In yeast, how metabolic signals control the progressive choreography of structural reorganizations observed in quiescent cells during a natural life cycle remains unclear. We have developed an integrated microfluidic device to address this question, enabling continuous single-cell tracking in a batch culture experiencing unperturbed nutrient exhaustion to unravel the coordination between metabolic and structural transitions within cells. Our technique reveals an abrupt fate divergence in the population, whereby a fraction of cells is unable to transition to respiratory metabolism and undergoes a reversible entry into a quiescence-like state leading to premature cell death. Further observations reveal that non-monotonous internal pH fluctuations in respiration-competent cells orchestrate the successive waves of protein super-assemblies formation that accompany the entry into a *bona fide* quiescent state. This ultimately leads to an abrupt cytosolic glass transition that occurs stochastically long after proliferation cessation. This new experimental framework provides a unique way to track single-cell fate dynamics over a long timescale in a population of cells that continuously modify their ecological niche.

## Introduction

Microorganisms have evolved plastic growth control mechanisms that ensure adaptation to dynamical environmental changes, including those that arise from their proliferation (such as nutrients limitations and cellular secretion in the medium). During its natural life cycle, budding yeast may undergo several metabolic transitions from fermentation to respiration, followed by entry into a reversible state of proliferation arrest known as quiescence (De Virgilio 2012; Gray et al. 2004; Sun and Gresham 2021; Miles, Bradley, and Breeden 2021). Despite quiescence being an essential part of the microorganism lifecycle that ensures cell survival over prolonged periods (Fontana, Partridge, and Longo 2010), it has received little attention compared to the analysis of biological processes in proliferative contexts.

Quiescent cells strongly differ from proliferating cells in terms of metabolic activity and gene expression (Gray et al. 2004; Miles et al. 2013). They also display a large body of structural rearrangements in the cytoskeleton, mitochondria, nuclear organization, and the appearance of protein super-assemblies, clusters, and aggregates (Sagot and Laporte 2019; Sun and Gresham 2021). So far, the complex and entangled regulatory processes controlling this particular state’s establishment remain poorly understood. In particular, the detailed sequence of events describing how the dynamics of metabolic cues during the natural lifecycle drive the entry into quiescence is still missing.

Also, an essential feature of microbial ecosystems in stationary phase is the existence of phenotypic variability (Campbell, Vowinckel, and Ralser 2016; Avery 2006; Holland et al. 2014; Labhsetwar et al. 2013; Ackermann 2015; Bagamery et al. 2020; Solopova et al. 2014) and history-dependent behaviors that lead to fate divergences (Balaban et al. 2004; Cerulus et al. 2018). In quiescence, the coexistence of heterogeneous cell populations has been previously reported (Allen et al. 2006; Laporte et al. 2018). Nevertheless, how phenotypic diversity emerges in a clonal population during a natural life cycle remains elusive(Fig. S1A). Bridging this gap requires performing longitudinal tracking of individual cells over time. However, an important technical obstacle is that it must be done in population-scale growth experiments to allow cell proliferation to have a collective impact on the environment.

Previous work has used an abrupt transition to glucose starvation to study how cells reorganize upon entry into stationary phase in various biological contexts(Munder et al. 2016; Bagamery et al. 2020). While this experimental framework may be helpful for studying standard properties of cells undergoing proliferation arrest, the results cannot be transposed to the context of entering quiescence during an undisturbed life cycle, in which cells undergo a sequence of metabolic transitions and continuously feed back into the composition of their environment. Indeed, an essential condition to reach a *bona fide* quiescence state (i.e. the ability to recover proliferation after prolonged arrest) is that cells must experience a respiration phase to accumulate carbohydrates which does not occur upon abrupt glucose starvation (Ocampo et al. 2012; Li et al. 2013). In addition, different nutrient limitations lead to distinct quiescent states (Klosinska et al. 2011). Therefore, it is essential to develop novel methods that capture the true dynamics of cell transitions as they may occur in their ecological niche(Miles, Bradley, and Breeden 2021).

Here, we report the development of a microfluidic platform for single-cell ecology, allowing continuous tracking of individual cells’ fate during an unperturbed full life cycle (up to 10 days). Using a fluorescent reporter of internal pH (Mouton et al. 2020; Miesenböck, De Angelis, and Rothman 1998) (Dechant et al. 2010; Munder et al. 2016), we observe that the diauxic shift witnesses a cell fate divergence, where a minority of cells experience a metabolic crash similar to that observed upon an abrupt starvation (Bagamery et al. 2020). Interestingly, our long-term tracking capabilities further reveal that these cells experience a premature yet reversible induction of cellular reorganizations that coincide with limited survival. In contrast, respiration-competent cells experience fluctuations in internal pH in sync with metabolic transitions that drive successive waves of cellular reorganizations and a stochastic switch to a glass transition of the cytoplasm long after proliferation cessation. Altogether, our analysis reveals how metabolic changes encountered by yeast cells during an unperturbed life cycle coordinate the temporal control of complex cellular reorganizations.

## Results

To track individual cell behavior during an unperturbed life cycle, we set up a device composed of a 25mL liquid yeast culture (YPD medium) connected to a microfluidic device for single-cell observation (Fig. 1A, Fig. S1B,C,F). Thanks to a close recirculation loop, individual cells trapped in the microfluidic device could be imaged over time while experiencing the same environmental changes as the liquid culture population. To prevent clogging in the microfluidic device due to the high cell density in the culture (up to 10^9^ cells/ml), we designed a filtration device based on inertial differential migration (Bhagat, Kuntaegowdanahalli, and Papautsky 2008). Using this technique, cells were rerouted back to the liquid culture before entering the microfluidic device (Fig. 1A). Optical density (O.D.) and fluorescence measurements revealed a filtration efficiency superior to 99%, reducing the concentration of cells entering the device by two orders of magnitude (Fig. S1E), and allowing us to image the cells over up to 10 days. We also checked that the same filtration device could sort *S. pombe* cells with 92% efficiency, thus highlighting the versatility of the methodology.

**Figure 1.**
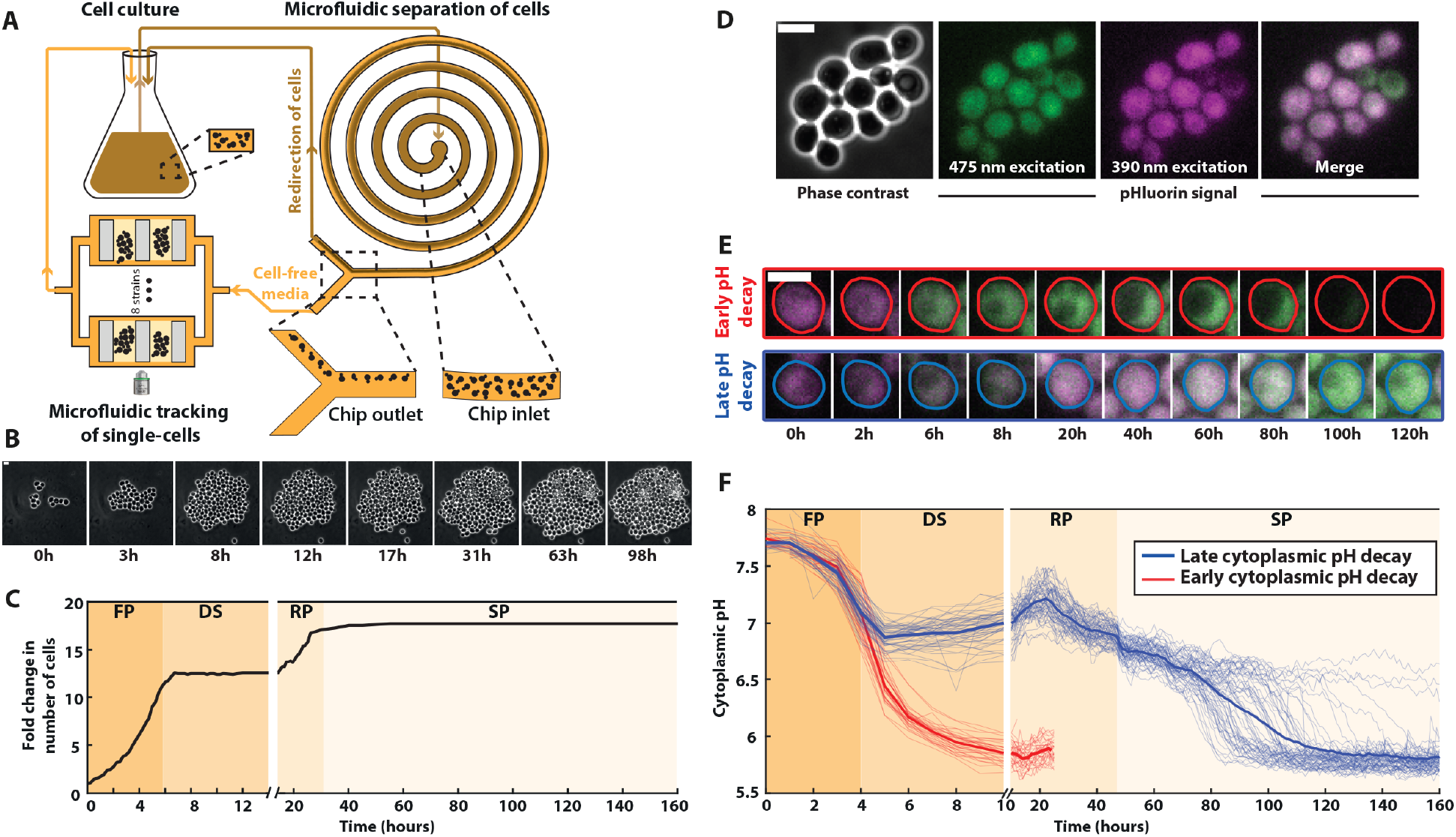
A microfluidic platform for single-cell tracking during the yeast proliferation cycle. A) Schematics of the experimental setup, representing the liquid culture flask, the observation microfluidic device with trapped cells, and the microfluidic filtering device designed to redirect the cells back to the liquid culture while recirculating the medium of the liquid culture to the observation microfluidic chamber. B) Sequence of phase-contrast images of cells growing in the microfluidic device.; Scale bar = 5 microns. C) Fold increase in cell number over an entire life cycle for the microcolony displayed in B; each shaded area represents a distinct proliferation phase, which was determined using piecewise linear fitting to cell proliferation data (see Methods and S1G for details): fermentation (FP), diauxic-shift (DS), respiration (RP), Stationary Phase (SP). D) Cluster of cells showing typical phase contrast, fluorescence, and overlay images using the cytoplasmic pH sensor pHluorin. E) Typical sequences of overlaid fluorescence images obtained with the pHluorin sensor at indicated time points. Colored lines indicate cell contours. F) Quantification of the absolute cytoplasmic pH as a function of time; each line represents an individual cell, while the bold line indicates the average among cells with either an early (red lines, N=32) or late (blue line, N=64) decaying pH.

Using this technique, we successfully recapitulated with single-cell resolution the successive proliferation phases occurring at the population level upon carbon source exhaustion (Richards 1928), namely: a rapid exponential growth (doubling time = 84 min +/−12 min) corresponding to glucose fermentation (referred to as the fermentation phase or FP in the following, from t=0 to t=5.5 h), followed by a sharp growth arrest, or diauxic shift (DS, from t = 5.5 h to t = 13.9 h); then, the resumption of a slow proliferative regime (doubling-time = 307 min +/− 52 min) which is associated with the use of ethanol as a carbon source for a respiratory metabolism (respiration phase, or RP, from t = 13.9 h to t = 31.6 h) and a final cell proliferation cessation occurring upon carbon source exhaustion, leading to stationary phase (SP), see Fig. 1B-C and Movie 1). The transition times between each metabolic phase were determined using piecewise exponential fits (Fig. S1G). By refeeding the cells with fresh YPD medium after ten days, we observed that up to ~80% of them re-entered the cell cycle within 5h (Fig. S1H). This result confirmed the reversibility of cell proliferation arrest and testified that cells establish *bona fide* quiescence in our growth conditions (Laporte et al. 2017).

A drop in medium pH has long been reported to coincide with the resources’ exhaustion during microbial growth (Burtner et al. 2009). Yet, how internal pH evolves over an entire lifecycle has never been investigated. To address this, we used the ratiometric fluorescent probe of cytoplasmic pH pHluorin (Fig. 1D, (Miesenböck, De Angelis, and Rothman 1998; Mouton et al. 2020)), which we calibrated to display the actual internal pH (Fig. 1D-F and Fig. S1I). Using this readout, we observed that the pH, which was initially around 7.7, started to decline synchronously in all cells during the F phase (Fig. 1E-F and Movie 2). At the onset of the DS, most cells (88%, N=466) abruptly reached a plateau (pH ~ 6.9, blue lines on Fig. 1F) followed by a slight pH increase (up to pH~7.2) that coincided with entry into a respiratory metabolism. In contrast, a minority (12%, N=466) of cells (Fig. 1E and red lines on Fig. 1F) experienced a further drop in pH down to about 5.8 during the DS. In this subpopulation, the fluorescence signal progressively disappeared, precluding monitoring the internal pH for more than 20h in a reliable manner.

In cells with high internal pH during the DS, the pH gradually declined after reaching a local maximum during the R phase. These cells then experienced a sharp pH drop down to about 5.8, which occurred at very heterogeneous times during SP, unlike the cells with an early pH drop. Altogether, these observations revealed unprecedented dynamics of internal pH during the yeast life cycle: pH variations appeared to be in sync with the sequence of proliferation phases, suggesting that internal pH is a crucial marker of the cells’ metabolic status during its lifecycle.

These continuous pH measurements also unraveled a divergence in cell fate at the DS, leading to the early emergence of heterogeneity within the population, in line with previous observations made upon abrupt starvation (Bagamery et al. 2020). Recent studies have shown that the activation of respiration was a crucial metabolic response to survive glucose deprivation (Weber et al. 2020), enabling carbohydrate storage (Ocampo et al. 2012) and long-term viability (Laporte et al. 2018). Interestingly, it was shown that a respiration defect was naturally observed in about 10% of cells upon proliferation cessation following glucose exhaustion (Laporte et al. 2018). Hence to further characterize whether differences in metabolic status drove the emergence of divergent cell fates at the DS, first, we quantified cellular proliferation over time using single-cell volume measurements. We found that cells with a late pH drop resumed growth and roughly doubled their biomass during the R phase (blue lines on Fig. 2A) in agreement Fig. 1C.

**Figure 2.**
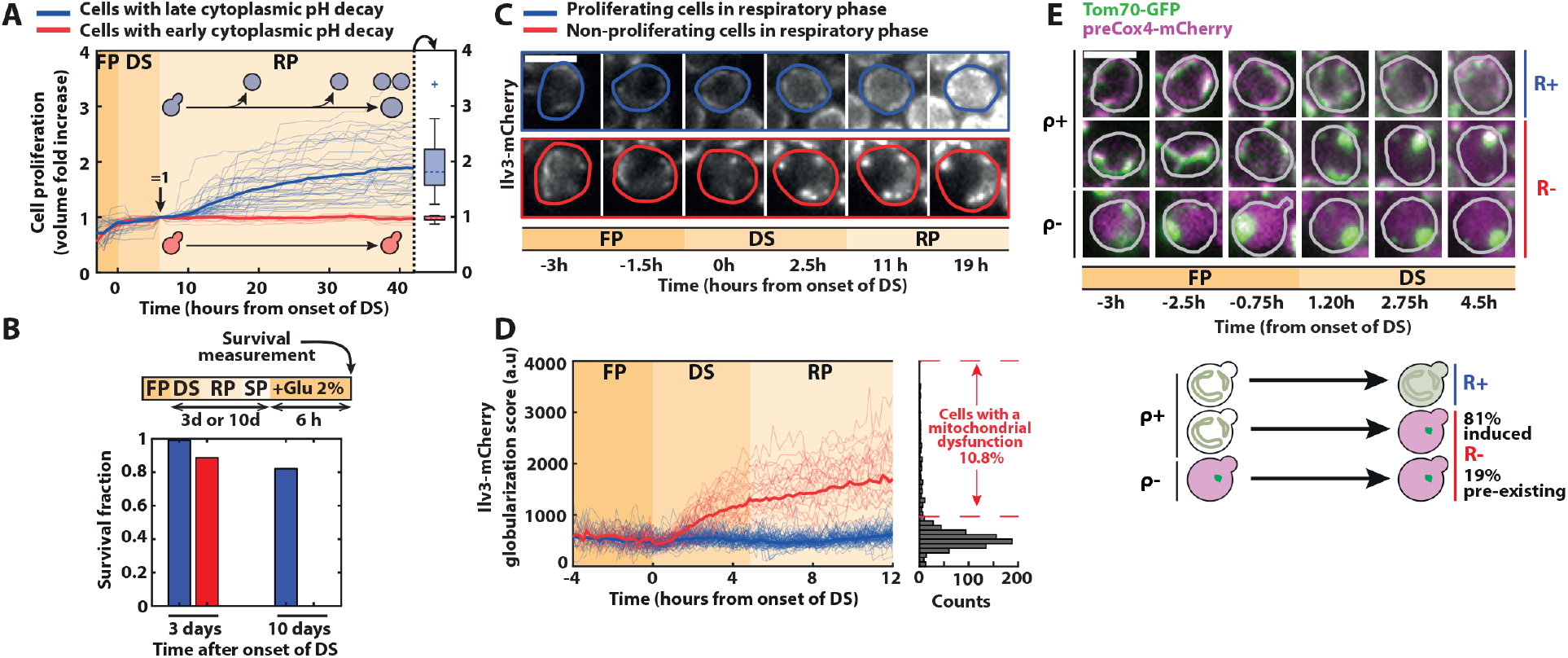
Divergent cell fates induced by a metabolic challenge at the DS. A) Quantification of single-cell growth during F, DS, and R phases, as defined in Fig. 1. Each line represents the fold area increase (including buds) of single cells over time, normalized by cell area at the end of the DS (N=50). The bold lines represent the averages over all the cells that experience fast (red) and slow (blue) pH decay, respectively. Right: box plot indicating the fold increase in cell volume in each subpopulation during the R phase (N=40 for the slow pH decay population, N=10 for the fast pH decay population). B) Fraction of surviving cells among adapting (blue bars) and non-adapting (red bars) cells, measured by quantifying the cells’ ability to resume growth 6h after re-introduction of fresh medium (2% glucose) at 3 (N=53 for red bars, N=221 for blue bar) or 10 days (N=114 for red bars, N=403 for blue bar) after the DS. Scale bar = 5 μm. C) Representative sequence of fluorescence images obtained with the Ilv3-mCherry mitochondrial marker at the indicated time points for the different classes of cells, as indicated by the colored contour. D) Single-cell quantification of a globularization score (see Methods) from fluorescence images over time for both adapting (blue line) (N=81) and non-adapting cells (red line) (N=28). The bold lines represent averages within each subpopulation. Right: histogram of globularization score for each cell (N=720). E) Pre-existing versus newly occurring respiratory defects in cells experiencing the DS; Sequence of fluorescent images (overlay of preCox4-mCherry and Tom70-GFP) at indicated times. Scale bar = 5 μm. Each line represents a different type of cell fates (top: rho+ R+; middle: rho+ R−; Bottom: rho− R−). Schematics: representation of the different cell fates based on the fluorescence patterns of preCox4-mCherry and Tom70-GFP, with the quantification of the fraction of each subpopulation (N=701).

In contrast, cells that experienced an early pH drop (red lines on Fig. 2A) did not recover during the R phase, suggesting that they could not transition to respiratory metabolism. Indeed, these cells did not resume growth when adding lactate (i.e., a non-fermentable carbon source) to the medium (Fig. S2A). However, most of them were still viable three days in SP - even though their survival declined faster than adapting cells (Fig. 2B).

Second, to assess cellular respiratory function, we used the mitochondrial Ilv3-mCherry marker (Laporte et al. 2018). Whereas the mitochondrial network architecture appeared similar (i.e., tubular) in all cells before the DS, proliferating cells in the R phase displayed a fragmented mitochondria phenotype typical of respiring cells (blue cell, Fig. 2C). In contrast, non-proliferating cells underwent a globularization of their mitochondrial network (red cell, Fig. 2C and Movie 3). We further quantified the mitochondrial network’s reorganization dynamics by computing a custom aggregation index that discriminates globularized versus tubular and fragmented mitochondria (Fig. 2D, see Methods for details). Based on the clear distinction in the aggregation index between adapting and non-adapting cells, we measured that about ~10 % of the cells could not transition to a respiratory metabolism (at t=12 h post-DS, Fig. 2D), in agreement with previous findings (Laporte et al. 2018). Importantly, this quantification revealed that the mitochondrial globularization in non-adapting cells was temporally closely associated with the DS since it started as early as 1h20 (p < 0.05) after its onset (Fig. 2C-D). Altogether, these results demonstrate that proliferating and non-proliferating cells experienced divergent cell fates at the DS based on their ability to switch to respiration; hence, they were referred to as respiration positive (R+) and negative (R−), respectively. Also, these results suggest that the inability of R-cells to activate a respiratory metabolism was either triggered by this metabolic challenge or, alternatively, pre-existed the DS, knowing that respiratory deficient cells (i.e., ρ-cells) are common in the BY background due to the genetic instability of mitochondrial DNA (Dimitrov et al. 2009).

To discriminate between these two hypotheses, we used the mitochondrial localization marker Tom70-GFP (Tom70 is a protein of the outer mitochondrial membrane) and a preCox4-mCherry fusion (preCox4 is a nuclear-encoded mitochondrial protein that is imported only in functional mitochondria (Veatch et al. 2009)), to assess the cells’ ability to respire (Fehrmann et al. 2013). Using these markers, we first checked that all R+ cells maintained a functional preCox4-mCherry import from fermentation to respiration (i.e., they were ρ+ cells, Fig. 2E). Then, we observed that among the R-cells, only a minority (i.e., 19%) had a dysfunctional mitochondrial import before the DS, indicating that they had a pre-existing respiratory deficiency (see ρ-cells on Fig. 2E). In contrast, the vast majority of R-cells (81%) transitioned from ρ+ to ρ-during the DS. This result demonstrates that, in contrast to a pre-existing condition, the respiration defect observed in R-cells occurred concomitantly with the environmental switch. Also, we showed that the progeny quite faithfully inherited it (Inheritance Index = 0.52, i.e., is smaller than 1, see Fig. S2B and Material and Methods for details). Altogether, these observations supported a scenario in which a fate divergence occurred early at the onset of the DS, where R+ cells quickly switched to a respiratory metabolism, while R-cells failed to do so.

As numerous cellular reorganizations occur during entry into quiescence (Sagot and Laporte 2019), we sought to quantitatively determine how they are coordinated with metabolic transitions during the lifecycle. Indeed, the metabolism controls internal cellular pH, which is known to induce significant physicochemical changes in the proteome, such as protein aggregation and phase transition (Munder et al. 2016; Dechant et al. 2010). To do this, we monitored the dynamics of formation of supra-molecular bodies associated with quiescence or the response to starvation: P-bodies formation (using the Dhh1-GFP fusion (Beckham et al. 2008; Balagopal and Parker 2009; Mugler et al. 2016)), metabolic or regulatory enzymes prone to aggregation (Gln1-GFP and Cdc28-GFP) (Narayanaswamy et al. 2009; Petrovska et al. 2014; Shah et al. 2014), actin bodies formation (Abp1-GFP)(Sagot et al. 2006), and proteasome storage granules (PSG, using the Scl1-GFP fusion)(Laporte et al. 2008; Peters et al. 2013). Indeed, these proteins have been described to form specific structures previously.

We found that the cellular reorganizations were highly coordinated with the sequence of metabolic phases and the cellular proliferation status (Fig. 3A): in R+ cells, Dhh1-GFP and Gln1-GFP (Movie 4) foci appeared at the onset of the DS, then were partially dissolved during the R phase, and reappeared upon entry into SP. Other hallmarks of proliferation cessation, such as actin bodies, PSGs, and other protein foci (Cdc28-GFP), showed up only at the end of the R phase. Importantly, we observed that R-cells also experienced a consistent formation of fluorescent foci for the markers that we monitored, yet, unlike R+ cells, they all appeared during the DS. In addition, surprisingly, all these foci ultimately disappeared, presumably as a consequence of the premature cell death observed in this subpopulation.

**Figure 3.**
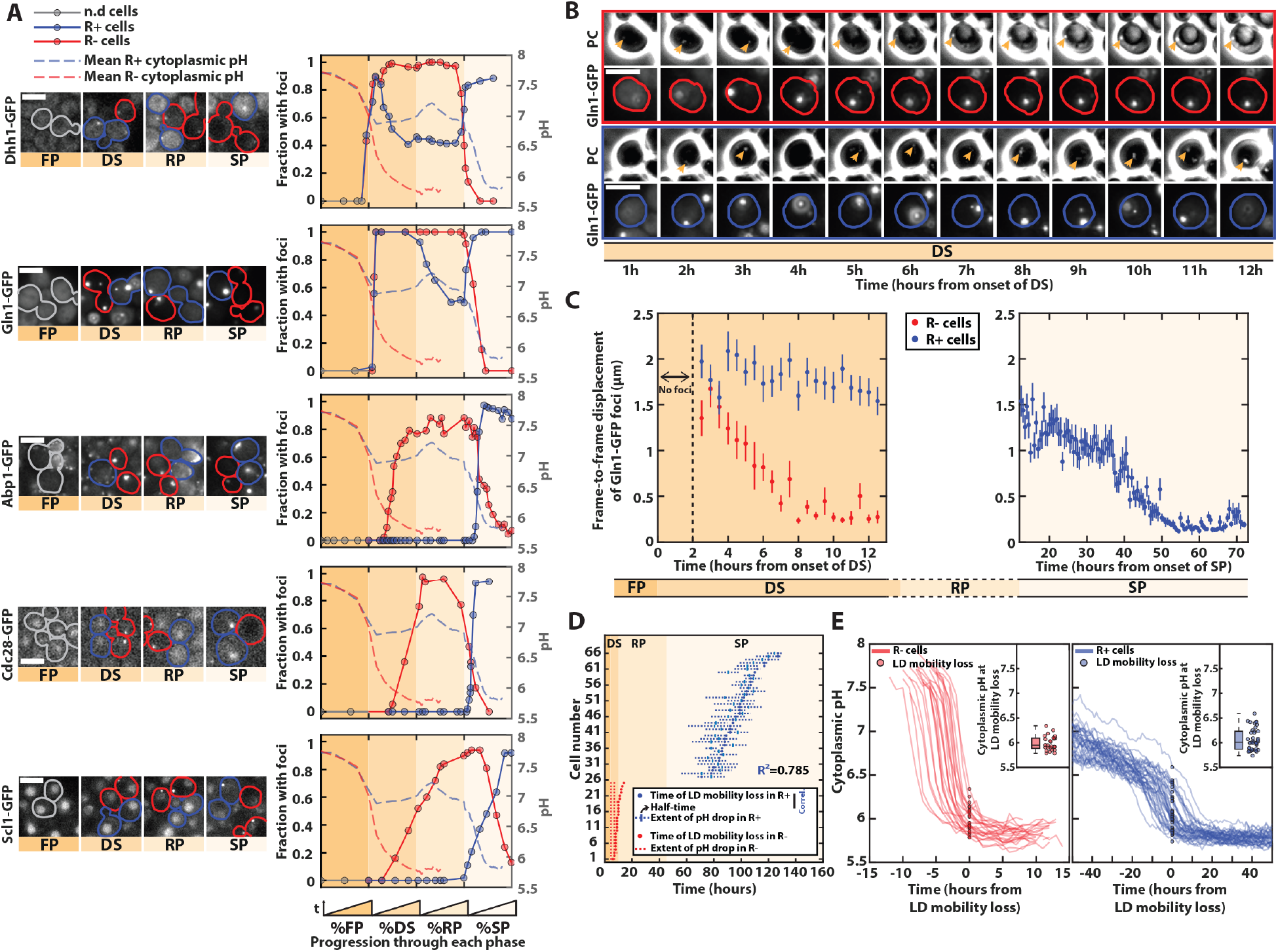
pH-driven phase-transition to a gel-like state upon proliferation cessation. A) Observation and quantification of fluorescent foci formation for the indicated fusion proteins (Dhh1-GFP, Gln1-GFP, Abp1-GFP, Cdc28-GFP, Scl1-GFP). Left: Each strip of fluorescence images displays unrelated cells at different phases during entry into SP; Colored contours indicate cells of interest (red for R-cells, blue for R+ cells, grey before the DS); Right: quantification of the fraction of cells with foci for each indicated fluorescent marker, as a function of the normalized time spent in each phase. Each solid colored line represents an indicated subpopulation of cells; The dashed colored lines represent the evolution of pH over time, based on data obtained in Figure 1; N>25 cells for each marker. Scale bar = 5 μm. B) Mobility of Gln1-GFP fluorescent foci and observation of lipid droplets (LD); Sequence of phase contrast and Gln1-GFP fluorescence images at indicated time points. The colored contours indicate the cells of interest (red and blue for R- and R+ cells, respectively). The orange arrowheads on the phase contrast images indicate the lipid droplet. Scale bar = 5 μm. C) Mobility of Gln1-GFP fluorescent foci; Quantification of average frame-by-frame displacement of Gln1-GFP foci for R+ and R-cells (blue and red points, respectively) starting after the appearance of foci (t > 2 hours after the onset of DS); Error bars represent the standard-error-on-mean (N=51 for R- and N=110 for R+); The right plot only features R+ cells, since Gln1-GFP foci are no longer present in the SP phase in R-cells (N=28); D) Temporal link between the drop in internal pH and the time of freezing of LD (N=25 for R- and N=43 for R+); Each line corresponds to a single cell and represents the extent of pH drop (see Material and Methods for details). Half-times of these drops are represented by a small vertical bar. The time of LD freezing is displayed as a dot. Of note, 2 cells didn’t display a pH drop, nor an LD freezing, hence are not displayed on the plot. E) Measurement of internal pH at the time of LD freezing; Overlay of internal pH in single cells obtained after synchronizing all traces with respect to the time of LD freezing, for R- (left) and R+ cells (right); Each dot represents the pH at the time of freezing in each cell (N=25 for R- and N=43 for R+); Inset: boxplot showing the distribution of pH values of every single cell at the time of LD freezing.

By overlapping the dynamics of fluorescence foci formation with that of internal pH (from Fig 1F), we noticed that the formation of bodies overall coincided with variation in absolute pH level in both R+ and R-cells, even though it appeared at different timescales. This finding is compatible with the hypothesis that it is the drop in internal pH - which reflects a decrease in metabolic activity - that drives successive waves of body formation and the appearance of quiescence hallmarks. Indeed, upon the DS, a pH decrease down to ~7 would trigger the formation of Gln1 and Dhh1 foci in both R+ and R-cells. In R+ cells, the re-rise of the pH associated with proliferation resumption in the R phase would induce the disassembly of these structures until the pH reaches 7 again, after carbone source exhaustion in SP. In both R+ and R-, Actin Bodies, PSG and Cdc28 foci would form only when the intracellular pH reaches 6.

To further check this hypothesis, since energy depletion was shown to induce a global modification of the cytoplasm to a glassy state (Munder et al. 2016), we sought to observe these transitions by monitoring the frame-to-frame displacement of Gln1-GFP foci over time (Fig. 3B, (Munder et al. 2016)). This marker was chosen because foci are already present at the DS. We found that the mobility of Gln1-GFP foci decreased sharply in R-cells during the DS (Movie 4) and much later in R+ cells, consistently with the differences observed regarding the times of pH drops in both populations (Fig. 3C). To better establish the links between pH and mobility, we exploited the fact that lipid droplets (LD) could be conveniently observed using phase-contrast images (Heimlicher et al. 2019). Consistently, the time of loss of LDs mobility correlated very well with that of Gln1-GFP foci (Fig S3A). By quantifying both the time of loss of LD mobility and the dynamics of pH drop in single cells, we observed that both events were tightly correlated in all cells, despite the large cell-to-cell heterogeneity in the time of pH drop in R+ cells (Fig 3D and S3B). Also, after synchronizing all single-cell trajectories from the time of loss of LD mobility, we showed that the freezing of the cytoplasm occurred at a similar pH ~ 6 in both R+ and R-cells (Fig 3E). Therefore, these observations suggest that the internal pH, which displays a very dynamic behavior during the yeast life cycle, induces waves of cellular structural remodeling that ultimately triggers a global transition of the cytoplasm to a gel-like state. This model is further supported by the concomitance between the premature pH drop and the phase transition in respiratory deficient (R−) cells. Altogether, it suggests that, upon nutrients exhaustion, cells undergo stereotypical pH-dependent structural reorganizations, no matter their respiratory status. R-cells’ inability to switch to respiration, which ultimately compromises the long-term viability in this subpopulation (Ocampo et al. 2012; Weber et al. 2020), makes this transition precocious and explains the divergent cell fates in the population at the DS.

## Discussion

In this study, we have developed a new microfluidic platform that allows us to monitor a full cell proliferation cycle in liquid culture with single-cell resolution. Individual cell tracking and quantitative fluorescence measurements provide a unique dynamic assessment of the successive metabolic transitions from fermentation to the stationary phase observed in a liquid culture submitted to nutrients exhaustion. In contrast to most previous studies that used abrupt environmental switches to investigate the metabolic response to starvation, our methodology recapitulates the unperturbed dynamics of nutrients experienced by cells during a life cycle in laboratory conditions. We envision that this methodology could be further applied to other contexts in which collective cell behavior impacts the environment, which in turn shapes individual cellular response - e.g., metabolic oscillations (Tu et al. 2005), cooperative behaviors (Dal Co et al. 2020; Campbell et al. 2015).

Continuous monitoring of cell growth and mitochondrial markers revealed a clear divergence in cell fate, where a minority of them failed to establish a respiratory metabolism, in line with recent observations obtained using an abrupt medium change (Bagamery et al. 2020). The dramatic drop in internal pH upon proliferation cessation in R-cells at the DS suggests that the failure to transition to respiration induces a major loss of energy homeostasis in this subpopulation, that may in turn compromise long term viability (Ocampo et al. 2012; Weber et al. 2020). By and large, this phenomenon could not be explained by pre-existing phenotypic differences in the population during the fermentation phase (Bagamery et al. 2020) but rather appeared to be triggered by the metabolic challenge associated with the exhaustion of glucose.

Previous studies have unraveled how changes in cytosolic pH - including upon nutrient exhaustion - control protein supramolecular assemblies (Peters et al. 2013; Petrovska et al. 2014). Our analysis shows that non-monotonous fluctuations in pH level occur in sync with metabolic transitions during an unperturbed life cycle. Further, it reveals that successive pH drops are closely temporally related with the formation of many protein bodies and granules, despite the large cell-to-cell temporal variability associated with the onset of these events. Interestingly, since the structural reorganization of many protein complexes is one of the well-described hallmarks of quiescence (Sagot and Laporte 2019), our observations thus suggest that internal pH acts as a key controller that drives waves of structural changes at well-defined pH levels during entry into quiescence. That R-cells display foci formation similarly as the R+ cells provides further support to a model of stereotypical structural reorganizations driven by energy depletion and transduced by internal pH level.

In addition to monitoring the super-assembly of specific markers, we showed that the transition to SP is accompanied by a transition of the cytoplasm to a glassy-like phase, hence transposing previous observations (Parry et al. 2014; Joyner et al. 2016; Munder et al. 2016) to the context of an unperturbed lifecycle in budding yeast (Heimlicher et al. 2019). Importantly, the appearance of the glassy-like state occurs right at the DS in the R-cells and is significantly delayed and variable in time in R+ cells. We propose that the start of respiration upon glucose exhaustion prevents a precocious glass transition, which might be detrimental to cell viability if it occurs in an uncoordinated manner with other cellular reorganizations processes (e.g., energy storage)(Ocampo et al. 2012; Weber et al. 2020).

The abrupt and concomitant transition of all cells during the DS shows how the rapid evolution of the environment at this precise moment drives cell behavior in a deterministic manner, in the same way as during abrupt starvation(Bagamery et al. 2020; Munder et al. 2016). Conversely, the remarkable cell-to-cell temporal heterogeneity during the transition to a glassy state in R+ cells suggests that the cells have a developmental program whose progression is partly stochastic, i.e., not entirely determined by the external environment. This last observation further supports that it is impossible to follow the process of entry into quiescence faithfully by imposing the dynamics of environmental changes(Miles, Bradley, and Breeden 2021). Further studies using our methodology may allow us to discover how the succession of the different key steps of this developmental process contributes to establishing the specific physiological properties of quiescent cells. (e.g., long-term survival and stress tolerance).

## Methods

### Strains

All strains used in this study are congenic to S288C (see Supplementary Table 1 for details).

### Cell culture

Freshly thawed cells were grown overnight. In the morning, 2mL of the culture was inoculated into a 25mL flask containing fresh YPD medium. After 5h, 2mL of culture was used to load the cells into the microfluidic device, and the rest was used as circulating media for the experiment. This 5h delay was chosen so that cells only spend 2 to 3 divisions in a fermentation phase to limit the number of cells in the microfluidic device at the DS.

### Microfluidics and microfabrication

#### Microfabrication

Microfluidic chips were generated using custom-made microfluidic master molds. The master molds were made using standard soft-photolithography processes using SU-8 2025 photoresist (Microchem, USA). The designs were made on AutoCAD (available upon request) to produce chrome photomasks (jd-photodata, UK). The observation device was taken from a previous study (Goulev et al. 2017).

The mold of the dust filter chip (see below for details) was made by spin-coating a 25μm layer of SU-8 2025 photo-resist on a 3” wafer (Neyco, FRANCE) at 2700 rpm for 30sec. Then, we used a soft bake of 7min at 95°C on heating plates (VWR) followed by exposure to 365nm UVs at 160 mJ/cm^2^ with a mask aligner (UV-KUB3 Kloé^®^, FRANCE). Finally, a post-exposure bake identical to the soft bake was performed before development using SU-8 developer (Microchem, USA).

The mold for the spiral-shaped cell filter device (see below for details) was obtained by spinning SU-8 2025 at 1750 rpm to achieve a 50μm deposit. Bakes were 6 min long at 95°C and UV exposure was done at 180 mJ/cm2. A hard bake at 150°C for 15min was then performed to anneal potential cracks and to stabilize the resist.

Finally, the master molds were treated with chlorotrimethylsilane to passivate the surface.

#### Microfluidic chip fabrication

The microfluidic devices were fabricated by pouring polydimethylsiloxane (PDMS, Sylgard 184, Dow Chemical, USA) with its curing agent (10:1 mixing ratio) on the different molds. The chips were punched with a 1mm biopsy tool (Kai medical, Japan) and covalently bound to a 24 × 50 mm coverslip using plasma surface activation (Diener, Germany). The assembled chips were then baked for 1 hour at 60°C to consolidate covalent bonds between glass and PDMS. Then, the dust filter chip was connected to the spiral cell filter which was in turn connected to the observation chip. All the connections used 1mm (outside diameter) PTFE tubes (Adtech Polymer Engineering, UK).

All medium flows were driven using a peristaltic pump (Ismatec, Switzerland) at a 100μL/min rate. The system was connected to the tank of media and cells described in the previous section. The observation chip was loaded with the cells of interest using a 5mL syringe and a 23G needle. Last, we plugged the cell-outlet of the spiral and the observation chip outlet into the tank so the system is closed and without loss of media or cells.

#### Microfluidic cell filter and dust filter

The spiral-shaped microfluidic device (Bhagat, Kuntaegowdanahalli, and Papautsky 2008) was designed to filter out cells coming from the liquid culture to prevent clogging in the observation device (Fig. S1C). The dimensions of the cell-filter (i.e. a channel of 100μm width and 50μm micron height, defining a spiral of 5-loops separated by 900μm) were set to maximize the separation of haploid yeast cells, i.e., ~5μm particles, according to the following principle: particles in a spiral microfluidic channel with a rectangular section are submitted to several inertial forces that depend on their size and are either directed towards the center of the channel or the walls. Therefore, particles of similar diameter reach an equilibrium position and tend to focus on a single line, allowing their separation from the rest of the fluid by splitting the output channel in two different outlets. A similar filter could be used with other microorganisms by adapting its dimensions.

We also added a particle filter before the spiral to avoid any clogging of the spiral because of dust particles or debris (Fig. S1B).

To measure the filtration efficiency of the cell filter, the cell concentration of the inlet and the two outlets was measured at 4 different time points (0h, 24h, 48h, and 120h) using a turbidity measurement (O.D. 660nm, Fisherbrand). The filtration efficiency was equal to 99%, independently of the inlet cell concentration.

### Microscopy

For all experiments except pH measurements, cells in the observation device were imaged using an inverted widefield microscope (Zeiss Axio Observer Z1). Fluorescence illumination was achieved using LED lights (precisExcite, CoolLed) and the light was collected using a 63× (N.A. 1.4) objective and an EM-CCD Luca-R camera (Andor). Standard GFP and mCherry filters were used.

For experiments using the pHluorin cytosolic pH probe, a Nikon Ti-E microscope was used along with a LED light (Lumencor) fluorescence illumination system. The fluorescence was measured using two excitation wavelengths using a standard roGFP2 filter set (AHF, peak excitation wavelengths 390nm/18nm and 475/28nm, beamsplitter 495nm, emission filter 525/50 nm). Emitted light was collected using a 60x N.A. 1.4 objective and a CMOS camera (Hamamatsu Orca Flash 4.0).

We used motorized stages to follow up to 64 positions in parallel throughout the experiment. Single plane-images were acquired every 15 min, 30min, 60min or 240min depending on the phase of the culture (high sampling rate during FP versus lower acquisition frequency in SP) to limit photodamage.

### Image processing and data quantification

#### Calibration of the cytosolic pH probe

To calibrate the probe, 2 mL of exponentially growing culture (0.5 OD600) were centrifuged and resuspended in 200 μL calibration buffer (50 mM MES, 50 mM HEPES, 50 mM KCl, 50 mM NaCl, 200 mM NH4CH3CO2) at various pH from 5 to 8, and supplemented with 75 μM monensin, 10 μM nigericin, 10 mM 2-deoxyglucose, and 10 mM NaN3, as described previously (Mouton et al. 2020). The cells were incubated in this buffer for 30min and then imaged in the microfluidic device to perform a ratiometric fluorescence measurement (Fig. S1I).

#### Image processing

The raw images were processed using Matlab-based software (PhyloCell) and custom additional routines (Fehrmann et al. 2013; Paoletti et al. 2016). This software has a complete graphical interface for cell segmentation (based on a watershed algorithm), tracking (assignment cost minimization), and fluorescence signal quantification. The software can be downloaded online (http://charvin.igbmc.science/Resources).

#### Data replicates

All measurements reported in this study are based on at least 3 replicates.

#### Quantification of growth rate

The growth rate of the cells was computed using the evolution of the area of segmented cells (and buds) over time.

#### Identification of growth phases

To determine the limit between successive growth phases, we used a piecewise linear fit to the evolution of total number of cells with time (D’Errico 2017). The time point of the intersection between two pieces of the piecewise linear fit was defined as the time of transition between the two corresponding phases (See Fig. S1G).

#### Globularization score

The Ilv3-mCherry globularization score was measured in each cell by calculating the mean intensity of the 5 brightest pixels of the cell minus that of the other pixels, as previously described (Cai, Dalal, and Elowitz 2008).

#### Inheritance of the respiration defect

To quantify the inheritance of the respiration defect, we measured the average standard deviation of the respiration status (after assigning a value of 0 if the cells are unable to respire and 1 if they are respiration competent) within individual microcolonies of cells (microcolony size from 10 to 20 cells). We then obtained an “inheritance index” by normalizing this value to the standard deviation obtained after taking all cells into account (i.e., with no distinction about their belonging to a given microcolony). Hence this index equals 1 when the respiration defect appears at random in lineages and 0 if the transmission of the phenotype is fully inheritable.

#### Quantification of fluorescence foci

The fraction of cells displaying foci of GFP was manually scored at different time points using the ImageJ Cell Counter plugin.

#### Mobility of fluorescent foci

The fluorescent foci in each cell were detected using the centroid position of the 5 brightest pixels. The frame-to-frame displacement of the foci was computed by iterating this procedure over all the frames, and then averaged over a population of cells.

#### Time of foci mobility loss and lipid droplets

The time of mobility loss was determined by visual inspection of successive images due to the low signal-to-noise ratio in the image.

#### Extend of pH drop

The start and end of the pH drops were determined using a piecewise linear adjustment, similar to what was used to determine the transition between metabolic phases.

## Acknowledgments

We thank Sara Mouton and Liesbeth Veenhoff for discussions and for sharing the pHluorin strain ahead of their publication, Audrey Matifas for constant technical support throughout this work, Titus Franzmann for fruitful discussions, Sophie Quintin and Sandrine Morlot for careful reading of the manuscript. We thank Denis Fumagalli at the Mediaprep facility. This work was supported by the Fondation pour la Recherche Médicale (FRM, B.J, and G.C.), the Agence Nationale pour la Recherche (T.A. and G.C.), the grant ANR-10-LABX-0030-INRT, a French State fund managed by the Agence Nationale de la Recherche under the frame program Investissements d’Avenir ANR-10-IDEX-0002-02.

---

**Supplementary Figure 1.**
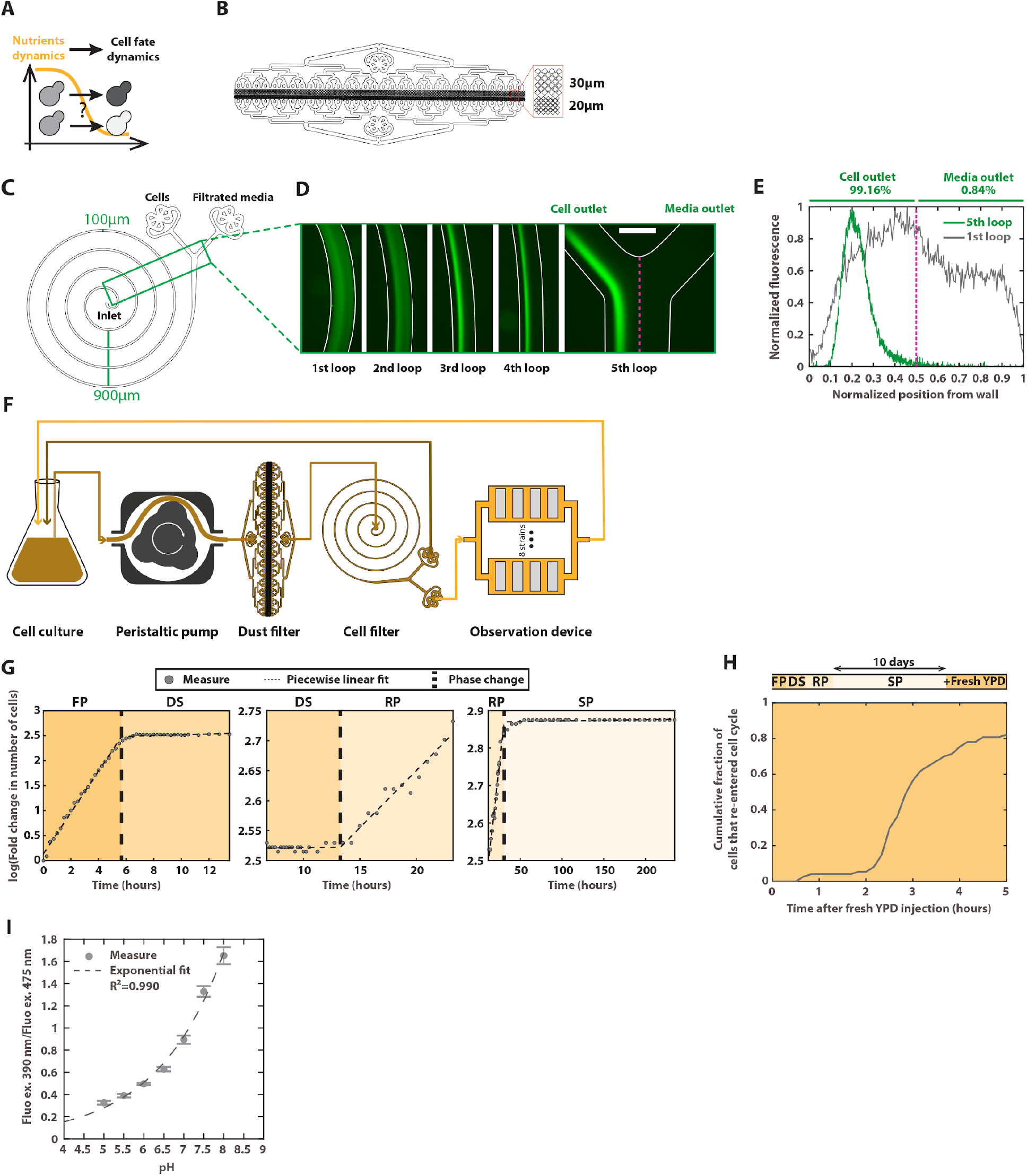
A) Schematics representing the main question addressed in this study, namely how the natural dynamics of environmental changes shapes non-uniform cellular responses and induces distinct cellular fates in a clonal population; B) Schematics of the dust filter device, including a close up of withe debris retention arrays of 30μm and 20μm size. C) Schematics of the cell-filtering device, made of a spiraling channel with 5 loops of 100μm width, separated by 900 μm. D) Sample fluorescence images of TDH3-GFP cells at different positions in the filtering device; Each image represents the indicated loop. The fushia dotted line represents the middle of the channel. Scale bar = 150μm. E) Fluorescence profil of the first loop (gray) and 5th loop (green) along the channel’s cross-section. The value is normalized to the maximum of each condition. In the upper part is displayed the cumulative signal in each side of the channel of the 5th loop. F) Schematics of the whole closed-loop fluidic platform; The flask is connected to a peristaltic pump, which drives media flows to the dust filter. The dust-free media then flows into the inlet of the spiral. The cell outlet of the spiral is redirected into the flask while the cell-free outlet irrigates the observation device. To close the loop, the outlet of the observation device is connected to the flask. G) Determination of the culture metabolic phases during the yeast life cycle based on the evolution of cell number over time; A piecewise linear fit on the number of cells defines the limit between FP and DS (left), DS and RP (middle) or RP and SP (right). Grey dots represent the measured number of cells in one microcolony, thin dashed lines represent the line fits and vertical dashed lines the transition time. Each shaded area represents a distinct proliferation phase, as indicated in the legend. H) Cumulative fraction of cells re-entering cell-cycle after 10 days in stationary phase over time, upon refeeding with YPD medium (N=60). I) Calibration curve of the pHluorin probe; Grey dots represent the ratio of fluorescence collected at 390nm over the fluorescence collected at 475 nm excitations, error bars represents the standard deviation (N=20); The dashed line correspond to an exponential fit f(x) = a*exp(b*x) of parameters (with 95% confidence bounds): a=0.01419 (0.005774, 0.0226) and b=0.5967 (0.5174, 0.6761).

**Supplementary Figure 2.**
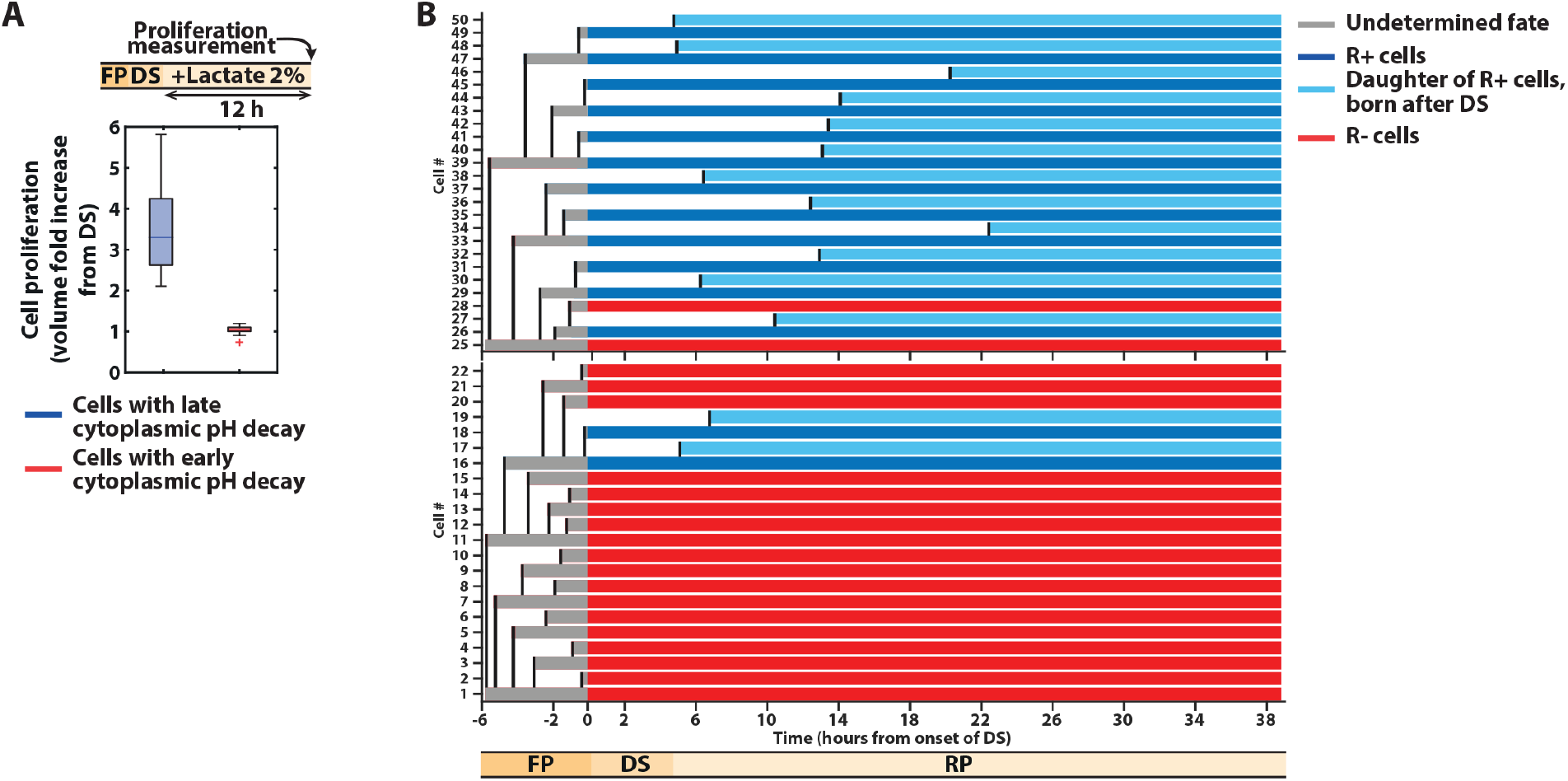
A) Proliferation capacity in a respirative media; Box plots indicate the fold increase in cell volume from the DS in each subpopulation (blue and red lines for cells with a late or early cytoplasmic pH decay, respectively), after a 12h exposure to a 2% lactate respirative media after the DS (N=30 for each box plot). B) Inheritance of the R+ and R-phenotype; Sample pedigrees representing 2 single cells (#1 and #25) and their progeny; each horizontal bar represents a cell. Cells reveal their respiratory status at the DS, hence are displayed in grey before the DS, and red (for R−) or blue (R+) after the DS. Daughters born after DS are represented in light blue. Only the first daughter of mothers born before the DS are represented for clarity.

**Supplementary Figure 3.**
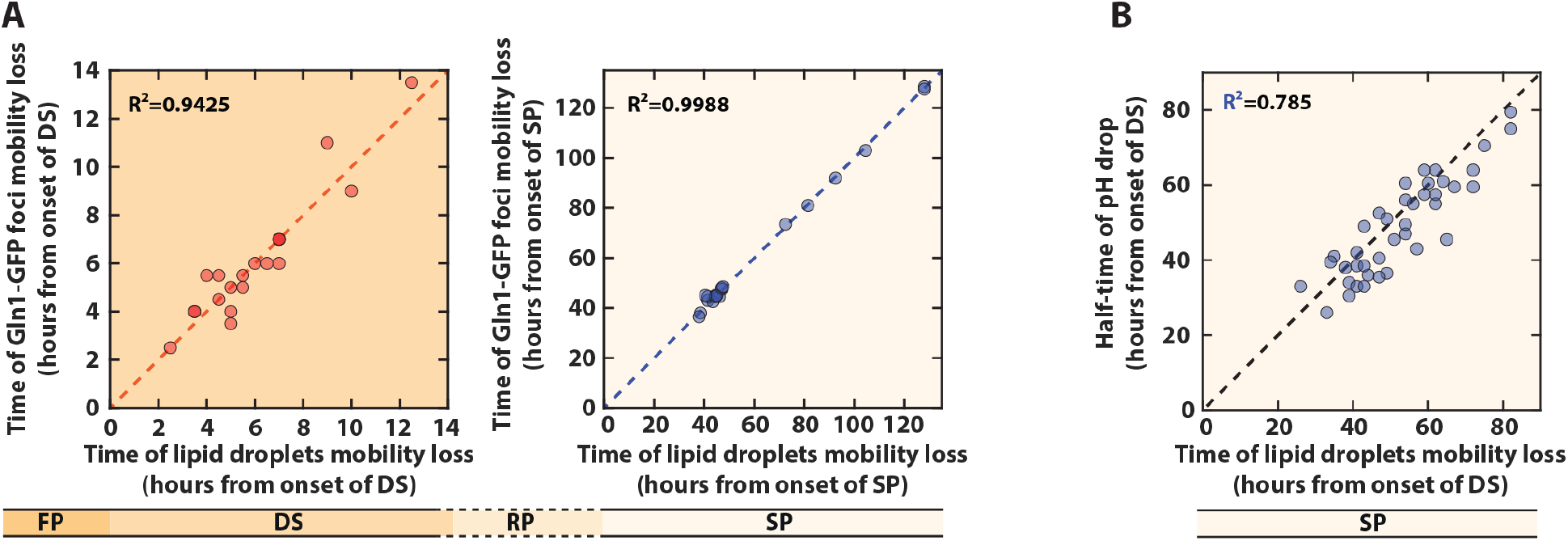
A) Correlation of the time of freezing of Gln1-GFP foci with that of the lipid droplet (LD) in R- (left plot) or R+ (right plot) cells; The origin of time is set as the proliferation arrest for each type of cells, i.e. the onset of DS and the onset of SP, respectively; The dashed lines represents the diagonal. B) Correlation of the half-time of the pH drop with the time of LD freezing in R+ cells; The time origin the onset of SP; The dashed lines represents the diagonal.

## Other supplemental data

### Supplementary Movie 1

Top: Phase contrast movie of a micro-colony growing in the observation device during the lifecycle of the culture. Bottom: Fold increase in the number of cells during colony proliferation. Colored progress bar indicates the metabolic phase of the culture as defined in Fig 1. Scale bar: 6.3μm

### Supplementary Movie 2

Top: phase contrast (left) and pHluorin (right, channel 1 and 2 merged as in Fig1) movie of a micro-colony growing in the observation device during the lifecycle of the culture. Colored progress bar indicates the metabolic phase of the culture as defined in Fig 1. The red and blue contours indicate a R- and a R+ cell, respectively. Scale bar: 10.8μm. Bottom: quantification of cytosilic pH over time, as described in Figure 1.

### Supplementary Movie 3

Phase contrast (left) and Ilv3-mCherry (right) movie of a micro-colony growing in the observation device during the lifecycle of the culture. Colored progress bar indicates the metabolic phase of the culture as defined in Fig 1. The thick red and blue contours indicate an original R- and a R+ cell, respectively while the thin red and blue contours indicate the daughters of the original cells. Scale bar: 6.3μm

### Supplementary Movie 4

Phase contrast (left) and Gln1-GFP (right) movie of a micro-colony growing in the observation device from fermentation phase to respiration phase. Colored progress bar indicates the metabolic phase of the culture as defined in Fig 1. The red and blue contours indicate a R- and a R+ cell, respectively. Scale bar: 6.3μm

**Supplementary Table 1.**
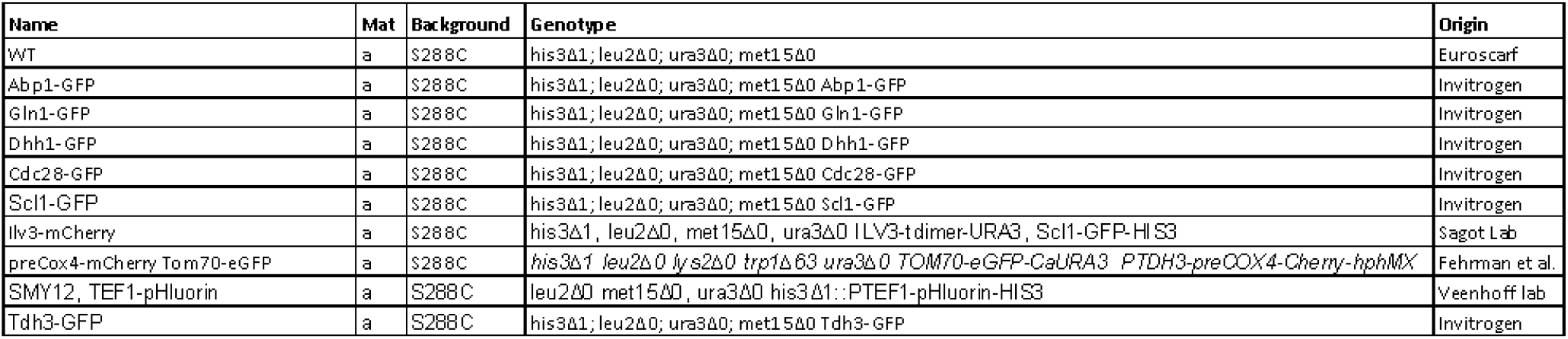
Strain Table

| Name | Mat | Background | Genotype | Origin |
| --- | --- | --- | --- | --- |
| WT | a | S288C | his3Δ1; leu2Δ0; ura3Δ0; met15Δ0 | Euroscarf |
| Abp1-GFP | a | S288C | his3Δ1; leu2Δ0; ura3Δ0; met15Δ0 Abp1-GFP | Invitrogen |
| Gln1-GFP | a | S288C | his3Δ1; leu2Δ0; ura3Δ0; met15Δ0 Gln1-GFP | Invitrogen |
| Dhh1-GFP | a | S288C | his3Δ1; leu2Δ0; ura3Δ0; met15Δ0 Dhh1-GFP | Invitrogen |
| Cdc28-GFP | a | S288C | his3Δ1; leu2Δ0; ura3Δ0; met15Δ0 Cdc28-GFP | Invitrogen |
| Scd1-GFP | a | S288C | his3Δ1; leu2Δ0; ura3Δ0; met15Δ0 Scd1-GFP | Invitrogen |
| Ilv3-mCherry | a | S288C | his3Δ1, leu2Δ0, met15Δ0, ura3Δ0 ILV3-tdimer-URA3, Scd1-GFP-HIS3 | Sagot Lab |
| preCox4-mCherry Tom70-eGFP | a | S288C | his3Δ1 leu2Δ0 lys2Δ0 trp1Δ63 ura3Δ0 TOM70-eGFP-CaURA3 PTDH3-preCOX4-Cherry-hphMX | Fehman et al. |
| SMY12, TEF1-pHluorin | a | S288C | leu2Δ0 met15Δ0, ura3Δ0 his3Δ1::PTEF1-pHluorin-HIS3 | Veenhoff lab |
| Tdh3-GFP | a | S288C | his3Δ1; leu2Δ0; ura3Δ0; met15Δ0 Tdh3-GFP | Invitrogen |

