## Supplementary figures and images for "Monitoring single-cell dynamics of entry into quiescence during an unperturbed lifecycle"

### Supplemental Figure 1

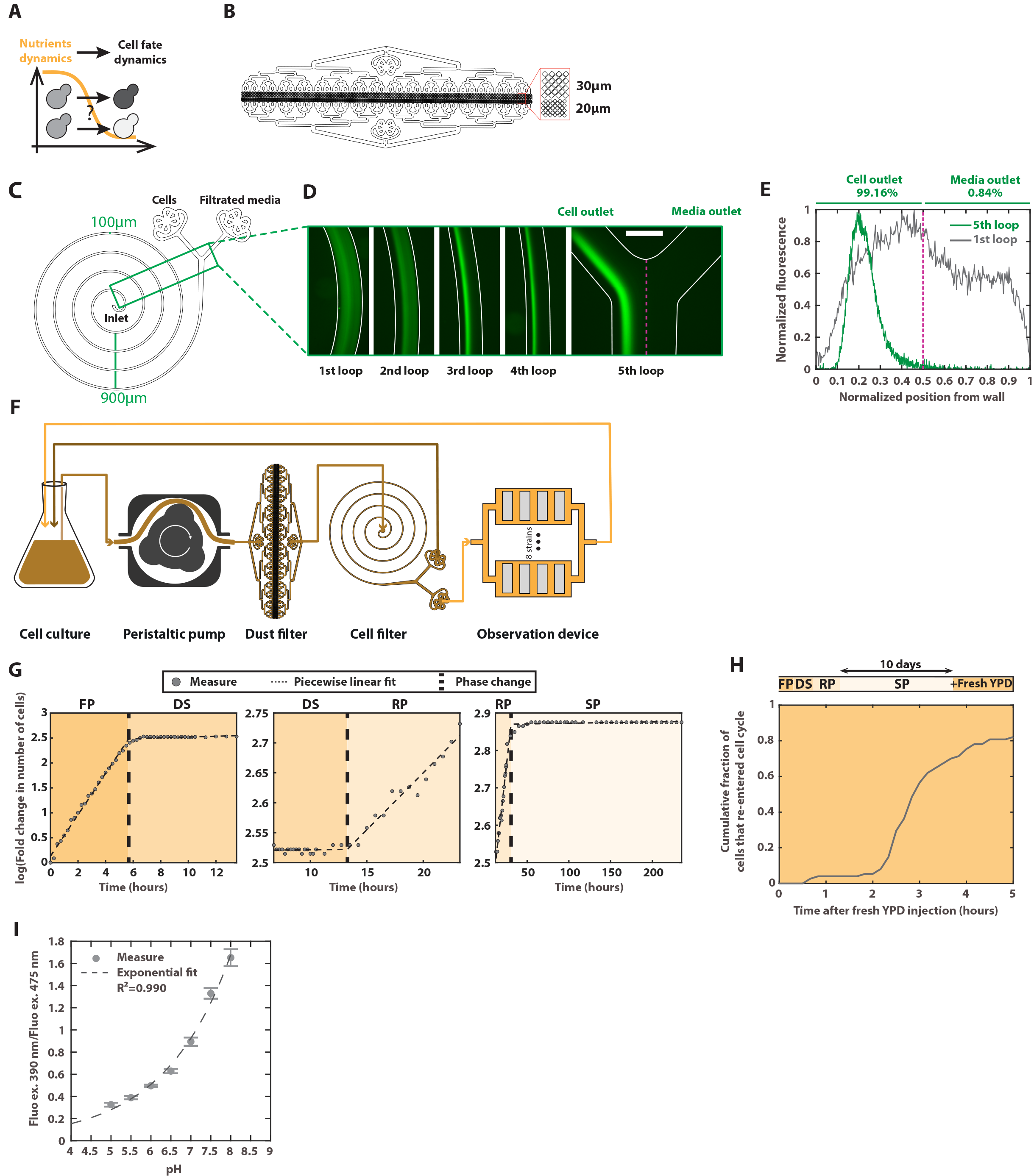

### Supplemental Figure 2

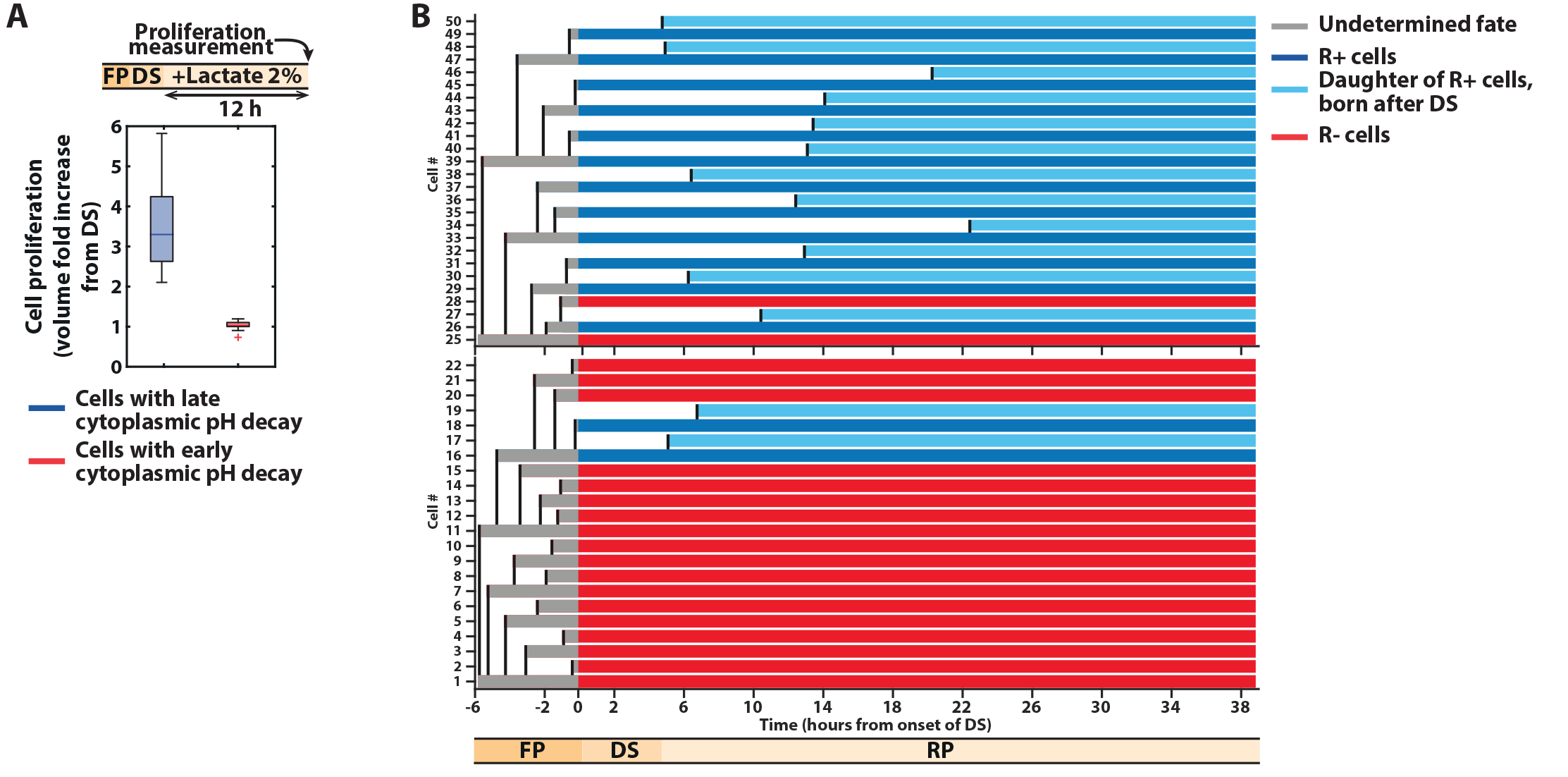

### Supplemental Figure 3

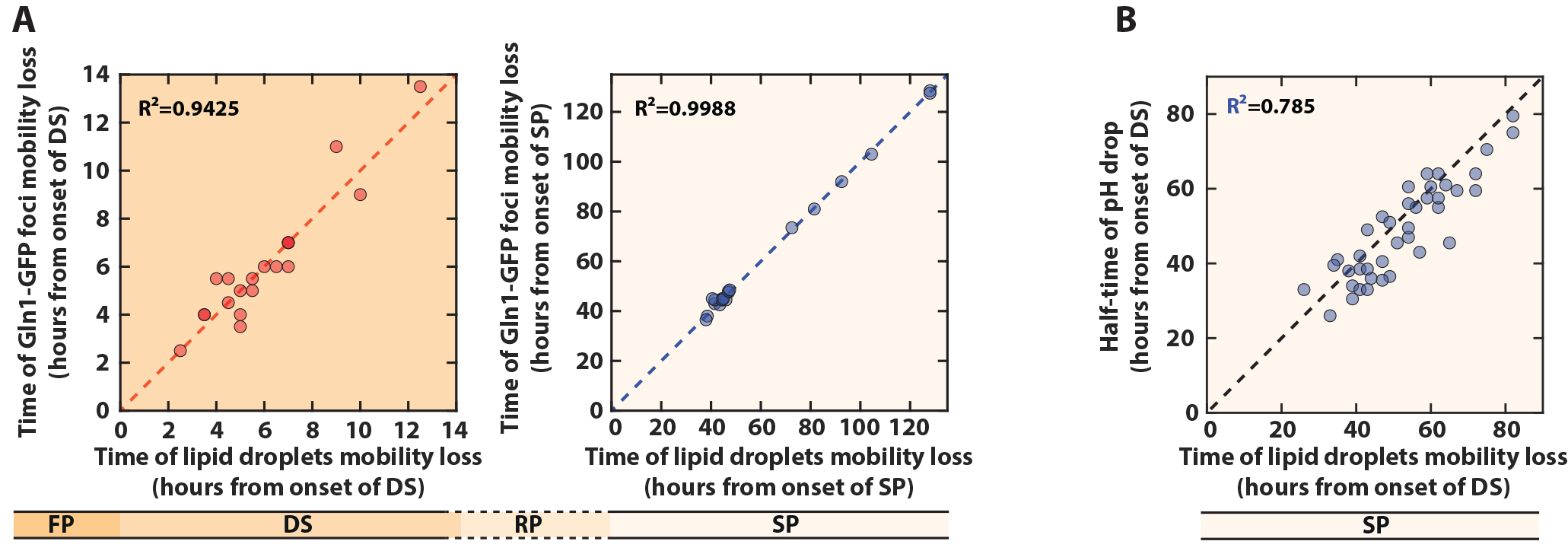
