## Supplemental Table 1 for "Monitoring single-cell dynamics of entry into quiescence during an unperturbed lifecycle"

| Name | Mat | Background | Genotype | Origin |
|---|---|---|---|---|
| WT | a | S288C | his3$\Delta$1; leu2$\Delta$0; ura3$\Delta$0; met15$\Delta$0 | Euroscarf |
| Abp1-GFP | a | S288C | his3$\Delta$1; leu2$\Delta$0; ura3$\Delta$0; met15$\Delta$0 Abp1-GFP | Invitrogen |
| Gln1-GFP | a | S288C | his3$\Delta$1; leu2$\Delta$0; ura3$\Delta$0; met15$\Delta$0 Gln1-GFP | Invitrogen |
| Dhh1-GFP | a | S288C | his3$\Delta$1; leu2$\Delta$0; ura3$\Delta$0; met15$\Delta$0 Dhh1-GFP | Invitrogen |
| Cdc28-GFP | a | S288C | his3$\Delta$1; leu2$\Delta$0; ura3$\Delta$0; met15$\Delta$0 Cdc28-GFP | Invitrogen |
| Scl1-GFP | a | S288C | his3$\Delta$1; leu2$\Delta$0; ura3$\Delta$0; met15$\Delta$0 Scl1-GFP | Invitrogen |
| Ilv3-mCherry | a | S288C | his3$\Delta$1, leu2$\Delta$0, met15$\Delta$0, ura3$\Delta$0 ILV3-tdimer-URA3, Scl1-GFP-HIS3 | Sagot Lab |
| preCox4-mCherry Tom70-eGFP | a | S288C | *his3$\Delta$1 leu2$\Delta$0 lys2$\Delta$0 trp1$\Delta$63 ura3$\Delta$0 TOM70-eGFP-CaURA3 PTDH3-preCOX4-Cherry-hphMX* | Fehrman et al. |
| SMY12, TEF1-pHluorin | a | S288C | leu2$\Delta$0 met15$\Delta$0, ura3$\Delta$0 his3$\Delta$1::PTEF1-pHluorin-HIS3 | Veenhoff lab |
| Tdh3-GFP | a | S288C | his3$\Delta$1; leu2$\Delta$0; ura3$\Delta$0; met15$\Delta$0 Tdh3-GFP | Invitrogen |
